## Supplemental materials for "The SARS-CoV-2 Omicron BA.1 spike G446S potentiates HLA-A*24:02-restricted T cell immunity"

### Extended Data Fig. 1

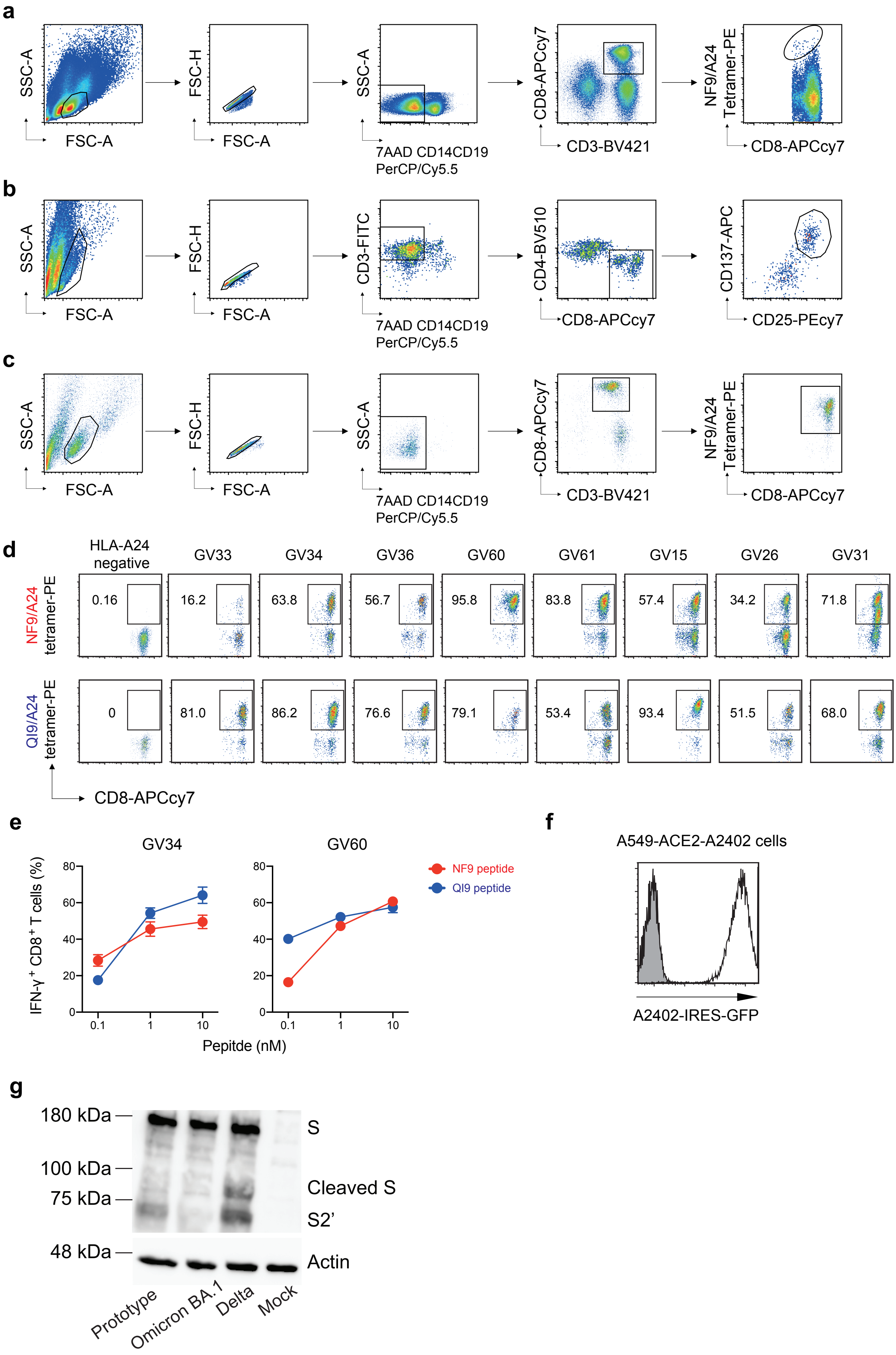

### Extended Data Fig. 2

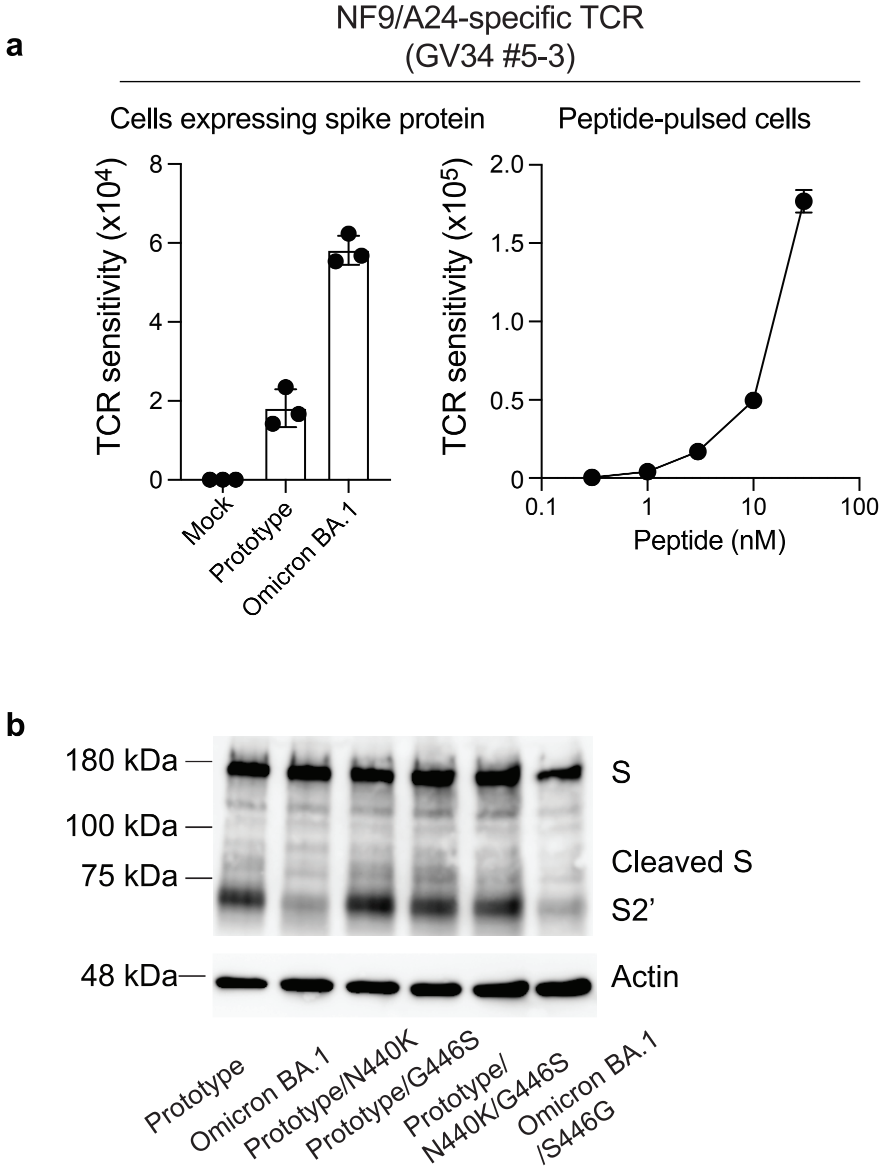

**Supplementary Table 1. Human PBMCs used in this study, related to Figure 1**

| Donor ID | Sex | Age | HLA-A24? | vaccinated? | Days after two doses of vaccination |
|---|---|---|---|---|---|
| Vku19 | Female | 36 | Positive | BNT162b2 | 21 |
| GV1 | Female | 28 | Positive | BNT162b2 | 9 |
| GV4 | Female | 18 | Negative | BNT162b2 | 20 |
| GV5 | Female | 18 | Positive | BNT162b2 | 20 |
| GV6 | Female | 79 | Positive | BNT162b2 | 21 |
| GV9 | Female | 24 | Positive | BNT162b2 | 21 |
| GV10 | Male | 23 | Positive | BNT162b2 | 22 |
| GV11 | Male | 23 | Positive | BNT162b2 | 22 |
| GV12 | Female | 28 | Negative | BNT162b2 | 23 |
| GV13 | Male | 22 | Positive | BNT162b2 | 21 |
| GV14 | Male | 23 | Positive | BNT162b2 | 22 |
| GV15 | Female | 23 | Positive | BNT162b2 | 21 |
| GV16 | Male | 22 | Positive | BNT162b2 | 22 |
| GV17 | Male | 24 | Negative | BNT162b2 | 21 |
| GV19 | Male | 24 | Positive | BNT162b2 | 22 |
| GV20 | Female | 24 | Positive | BNT162b2 | 22 |
| GV21 | Male | 22 | Positive | BNT162b2 | 21 |
| GV22 | Male | 22 | Positive | BNT162b2 | 22 |
| GV23 | Male | 25 | Positive | BNT162b2 | 22 |
| GV24 | Male | 23 | Positive | BNT162b2 | 22 |
| GV25 | Male | 24 | Negative | BNT162b2 | 22 |
| GV26 | Male | 23 | Positive | BNT162b2 | 22 |
| GV27 | Female | 23 | Negative | BNT162b2 | 21 |
| GV28 | Female | 23 | Positive | BNT162b2 | 21 |
| GV29 | Male | 23 | Positive | BNT162b2 | 21 |
| GV31 | Male | 23 | Positive | BNT162b2 | 22 |
| GV32 | Male | 56 | Positive | BNT162b2 | 27 |
| GV33 | Male | 39 | Positive | BNT162b2 | 24 |
| GV34 | Female | 38 | Positive | BNT162b2 | 24 |
| GV35 | Male | 52 | Positive | BNT162b2 | 24 |
| GV36 | Male | 39 | Positive | BNT162b2 | 24 |
| GV51 | Male | 34 | Positive | BNT162b2 | 30 |
| GV59 | Male | 37 | Positive | BNT162b2 | 126 |
| GV60 | Male | 51 | Positive | mRNA-1273 | 116 |
| GV61 | Female | 36 | Positive | mRNA-1273 | 107 |

**Supplementary Table 2. TCR sequences specific for the NF9/A24 and QI9/A24, related to Figure 3**

| Specificity | ID | TRAV | TRAJ | CDR3α | TRBV | TRBJ | TRBD | CDR3β |
|---|---|---|---|---|---|---|---|---|
| NF9/A24 | GV34 #2-2 | TRAV12-1*01 | TRAJ33*01 | CVVNALMDSNYQLIW | TRBV5-1*01 | TRBJ2-7*01 | TRBD1*01 | CASSLGQGYEQYF |
| | GV34 #5-3 | TRAV12-1*01 | TRAJ33*01 | CVVNLFDSNYQLIW | TRBV2*01 | TRBJ2-7*01 | TRBD1*01 | CASSEGAGYEQYF |
| | VKU19 #12-3 | TRAV12-3*01 | TRAJ44*01 | CAFTGTASKLTF | TRBV7-8*01 | TRBJ2-1*01 | TRBD2*02 | CASSPELNEQFF |
| QI9/A24 | GV33 #57 | TRAV3*01 | TRAJ8*01 | CAGVLFNTGFQKLVF | TRBV20-1*02 | TRBJ2-1*01 | TRBD2*01 | CSASDRGASGSFSNEQFF |
| | GV34 #43 | TRAV21*01 | TRAJ43*01 | CAAPRYNNNDMRF | TRBV2*01 | TRBJ2-2*01 | TRBD1*01 | CASSEGADAGELFF |
| | GV36 #10-2 | TRAV19*01 | TRAJ9*01 | CALSEPPSGGFKTIF | TRBV20-1*01 | TRBJ1-1*01 | TRBD1*01 | CSARGQGLNTEAFF |

**Supplementary Table 3. Primers for the construction of spike derivatives, related to Figures 3**

| Product | | Primer name | Sequence (5'-to-3') |
|---|---|---|---|
| Prototype spike | N440K | N440K Fwd | CAGCAACAAGCTGGACAGCAAGGTGG |
| | | N440K Rev | CTGTCCAGCTTGTTGCTGTTCCAGGC |
| | N440K G446S | N440K G446S Fwd | GCAACAAGCTGGACAGCAAGGTGTCCGGCAACTACAAC |
| | | N440K G446S Rev | GTTGCCGGACACCTTGCTGTCCAGCTTGTTGCTGTTC |
| | G446S | G446S Fwd | CAAGGTGTCCGGCAACTACAACTACCTC |
| | | G446S Rev | GTAGTTGCCGGACACCTTGCTGTCCAG |
| Omicron BA.1 spike | S446G | S446G Fwd | GACAGCAAGGTGGGAGGCAACTACA |
| | | S446G Rev | GCCTCCCACCTTGCTGTCCAGCTTG |
